# Inheritance of Chromatin Proteins in Budding Yeast: metabolic gene regulators TUP1, FPR4 and Rpd3L are retained in the mother cell

**DOI:** 10.1101/644138

**Authors:** Pauline Vasseur, Saphia Tonazzini, Francesc Rubert Castro, Iva Sučec, Arame Fall, Khadija El Koulali, Serge Urbach, Marta Radman-Livaja

**Affiliations:** Institut de Génétique Moléculaire de Montpellier, UMR 5535 CNRS, 1919 route de Mende, 34293 Montpellier cedex 5, France; Université de Montpellier, 163 rue Auguste Broussonnet, 34090 Montpellier, France.; CIRAD, Campus international de Baillarguet, UMR 117-ASTRE “Animal, Santé, Territoires, Risques & Écosystèmes”, TA A-117/E, Bât. G bureau 212, 34398 Montpellier cedex 5, France; IGF, CNRS, INSERM, Univ. Montpellier, F-34094 Montpellier, France

## Abstract

Asymmetric division is a prerequisite for cellular differentiation. Phenotypic transformation during differentiation is a poorly understood epigenetic phenomenon, in which chromatin theoretically plays a role. The assumption that chromatin components segregate asymmetrically in asymmetric divisions has however not been systematically tested. We have developed a live cell imaging method to measure how 18 chromatin proteins are inherited in asymmetric divisions of budding yeast. We show that abundant and moderately abundant maternal proteins segregate stochastically and symmetrically between the two cells with the exception of Rxt3, Fpr4 and Tup1, which are retained in the mother. Mother retention seems to be the norm for low abundance proteins with the exception of Sir2 and the linker histone H1. Our in vivo analysis of chromatin protein behavior in single cells highlights general trends in protein biology during the cell cycle such as coupled protein synthesis and decay, and a correlation between half-lives and cell cycle duration.

## Introduction

During asymmetric cell division a stem cell produces two different daughter cells: one cell that preserves the characteristics of “stemness” and will go on to perpetuate the stem cell lineage, and the other that will eventually undergo cellular differentiation into a specific cell type. Since the DNA sequence, barring random mutations introduced by DNA replication is identical in the two daughters, the stem cell phenotype and the differentiating phenotype are both inherited epigenetically: each cell will receive a different set of epigenetically encoded instructions, one to remain a stem cell and the other to start differentiating.

In order to understand how the two phenotypes are established and maintained we need to identify the epigenetic factors that distinguish the two daughters after asymmetric cell division.

Eukaryotic genomes are packaged into chromatin fibers consisting of arrays of nucleosomes: globular histone protein-DNA complexes^1^. Since chromatin structure together with chromatin bound proteins regulates transcription, it has the potential to transmit epigenetic information about the gene expression state of its underlying locus to the next generation (Petruk et al., 2012).

In order for chromatin features to be truly epigenetic, they have to be accurately “copied” (i.e. re-established on the correct genomic locus) after cell division and they have to be instructive of the transcription state at their genomic location. Since recent studies have provided experimental support for both claims -at least for some chromatin features such as heterochromatic histone marks-^2–4^, it is reasonable to assume that the daughter cell that inherits the maternal phenotype after asymmetric cell division should also inherit those parental chromatin components that presumably define that particular phenotype. Consequently these parental chromatin components would have to segregate asymmetrically only into one of the two cells, while the other daughter should inherit new chromatin components that will commit it to differentiation. On the other hand, chromatin features that are associated with constitutive traits that have to be maintained in both cells would be “inherited” symmetrically in both cells.

In order to understand cellular differentiation (pathological or developmental), it is important to identify which chromatin components carry epigenetic information and establish how these components segregate during genome replication and cellular division.

Budding yeast cells also divide asymmetrically, resulting in a larger mother and a smaller daughter cell. The mother can generate ~30 daughters during its replicative lifespan. The mother cell has therefore a phenotypic identity that distinguishes her from her daughters and that determines the length of her replicative lifespan. The ageing phenotype of the mother cell has been extensively studied and several models have been proposed to explain asymmetric segregation of ageing factors. Yet epigenetic factors that define the “mother” and “daughter” phenotypes and the role of chromatin in determining those phenotypes are still largely unknown. According to currently proposed models, the ageing phenotype is caused by molecular “ageing factors”. An ageing factor has to satisfy three criteria: it has to accumulate over time, it has to be preferentially retained in the mother cell and it has to directly or indirectly lead to cell death. Several candidates for “ageing factors” have been proposed, the most attractive being: 1) Extra chromosomal Ribosomal DNA Circles (ERCs)^5^, 2) protein carbonyls and 3) proteins damaged by oxidation, and old mitochondria (reviewed in^6^). Since none of these factors completely satisfy all the criteria, a consensus on the determining ageing agent has still not been reached, and epigenetic factors that define the “mother” phenotype and the role of chromatin in the process are still not known.

Several studies in recent years have identified proteins that accumulate asymmetrically in only one of the two cells after division. Yang e al.^7^ found 74 mother-enriched and 60 daughter-enriched proteins, however their assay could not differentiate between maternal proteins that potentially carry epigenetic information and newly synthesized copies. It is therefore unclear whether any epigenetic information is transmitted by chromatin associated proteins, which incidentally had a tendency to accumulate in the daughter cell in their assay. On the other hand, Thayer et al.^8^ looked specifically for long lived maternal proteins that accumulate in the aging mother cell. They mostly identified cytoplasmic and membrane proteins however, which probably don’t play a role in the epigenetic inheritance of gene expression states even though they could potentially contribute to the ageing phenotype of the mother cell, although the latter has not been explicitly tested. Finally, Garciá del Arco et al.^9^ show that during cell division of the fly midgut epithelium, the maternal CENP-A centromeric histone variant segregates asymmetrically into the daughter cell that will remain a stem cell.

Asymmetrically segregating chromatin components seem to be good candidates for “ageing factors” and epigenetic information carriers of mother or daughter phenotypes. We therefore set out to develop a live imaging screen aimed at identifying chromatin factors that could potentially be involved in epigenetic inheritance during asymmetric cell division. We used the photo-convertible fluorescent protein Dendra2. Since photo-conversion from green to red fluorescence after UV light irradiation is irreversible, one can readily measure the half-lives of maternal Dendra2 fusion proteins. Maternal proteins can be followed for several cell generations (one yeast cell generation is typically 90-100min long), unlike with a tandem sfGFP-mCherry construct that can only track newly synthesized proteins for 45min after synthesis due to the difference in folding kinetics of the fast folding sfGFP and the slow folding mCherry^10^.

We show that moderately and highly abundant maternal chromatin proteins segregate stochastically between mothers and their daughters with a cell population mean for the fraction of proteins retained in the mother at ~0.5 as expected. There are however several notable exceptions: Rxt3 (a subunit of the histone deacetylase complex Rpd3L), Fpr4 (proline isomerase of H3P38) and Tup1 (transcription repressor), which are preferentially retained in the mother cell. Surprisingly, low abundance proteins (between ~600 and ~3000 molecules per cell) are also mostly retained in the mother cell with the exception of Sir2 (part of the heterochromatic complex Sir) and H1 (linker histone), which segregate stochastically like more abundant proteins.

## Results

We have constructed a set of 18 strains carrying chromatin associated proteins fused to the photo-convertible fluorescent protein Dendra2 (Table S1). Dendra2 switches irreversibly from green to red fluorescence after a UV pulse. Consequently, live fluorescence imaging (HiLo,^11^) of dividing cells allows us to distinguish between already synthesized maternal proteins, which emit in the red spectrum after a UV pulse, and newly synthesized green proteins made after the pulse (**Figure 1**).

**Figure 1:**
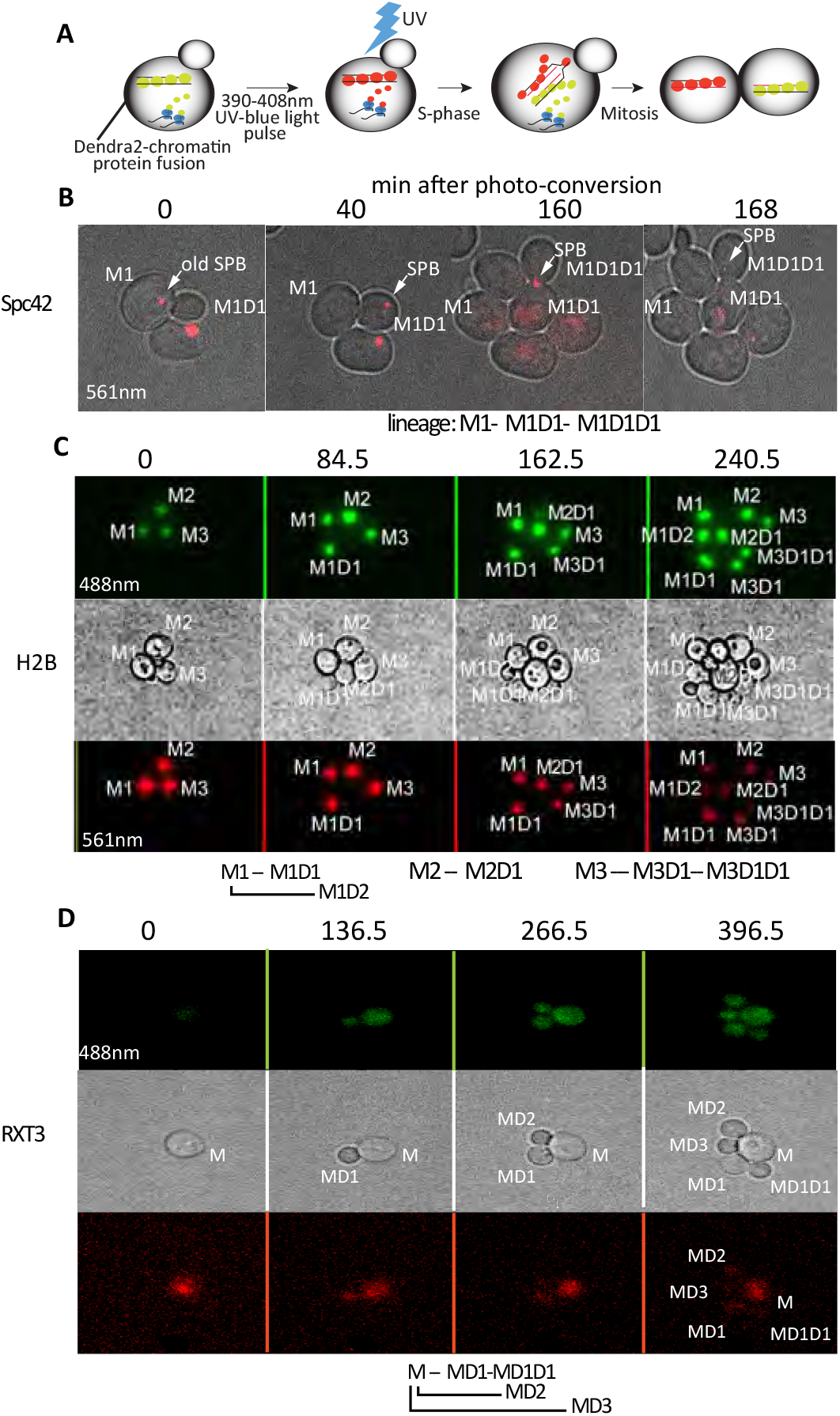
Tracking segregating maternal chromatic proteins. A. Chromatin proteins are fused to the photo convertible fluorescent protein dendra2. Proteins already incorporated in chromatin will switch from green to red fluorescence after UV light irradiation. Newly synthesized green fluorescent fusion proteins are incorporated into daughter chromatids after the UV pulse, and segregation of old red and new green proteins is monitored by live cell imaging. B-D HiLo imaging of Spc42(B) (component of SPB-Spindle Pole Body) fused to Dendra2. The “old” (red) SPB segregates into the daughter cell, as previously reported. Histone H2B (C) segregates symmetrically but can also be preferentially retained in the mother cell: compare red signals between M3D1 and its daughter at the 240.5min time point. Maternal RXT3 (subunit of the histone deacetylase Rpd3L) is retained in the mother cell (D). Cell lineages are shown below the images.

We can therefore record segregation patterns of maternal chromatin proteins, and follow cellular localization dynamics of fusion constructs through mitosis. Due to closed mitosis in yeast, nuclear localization during mitosis does not automatically signify chromatin association. Nevertheless, the inheritance of maternal proteins within the nucleus- as is the case for all tested proteins- is still an indication that these chromatin proteins may remain associated with mitotic chromosomes and could therefore be epigenetically inherited (for examples see **Figure 1 C**).

We used Spc42-dendra2 as a positive control. Spc42 is part of the Spindle Pole Body (SPB). Our analysis shows a preferential inheritance of the “old” SPB by the daughter cell as previously reported^10^ (**Figure 1B**). Our conditions for UV irradiation do not affect cell growth and viability as doubling times and the numbers of dividing cells are comparable between non-irradiated cells or cells grown in liquid media, and UV irradiated cells (Figure 3 and data not shown).

### Half-life measurement and protein segregation after cell division

The segregation patterns of maternal chromatin proteins between mother and daughter cells are estimated from the fraction of maternal proteins remaining in the mother cell after the first cell division following photo conversion (**Figure 2**).

**Figure 2:**
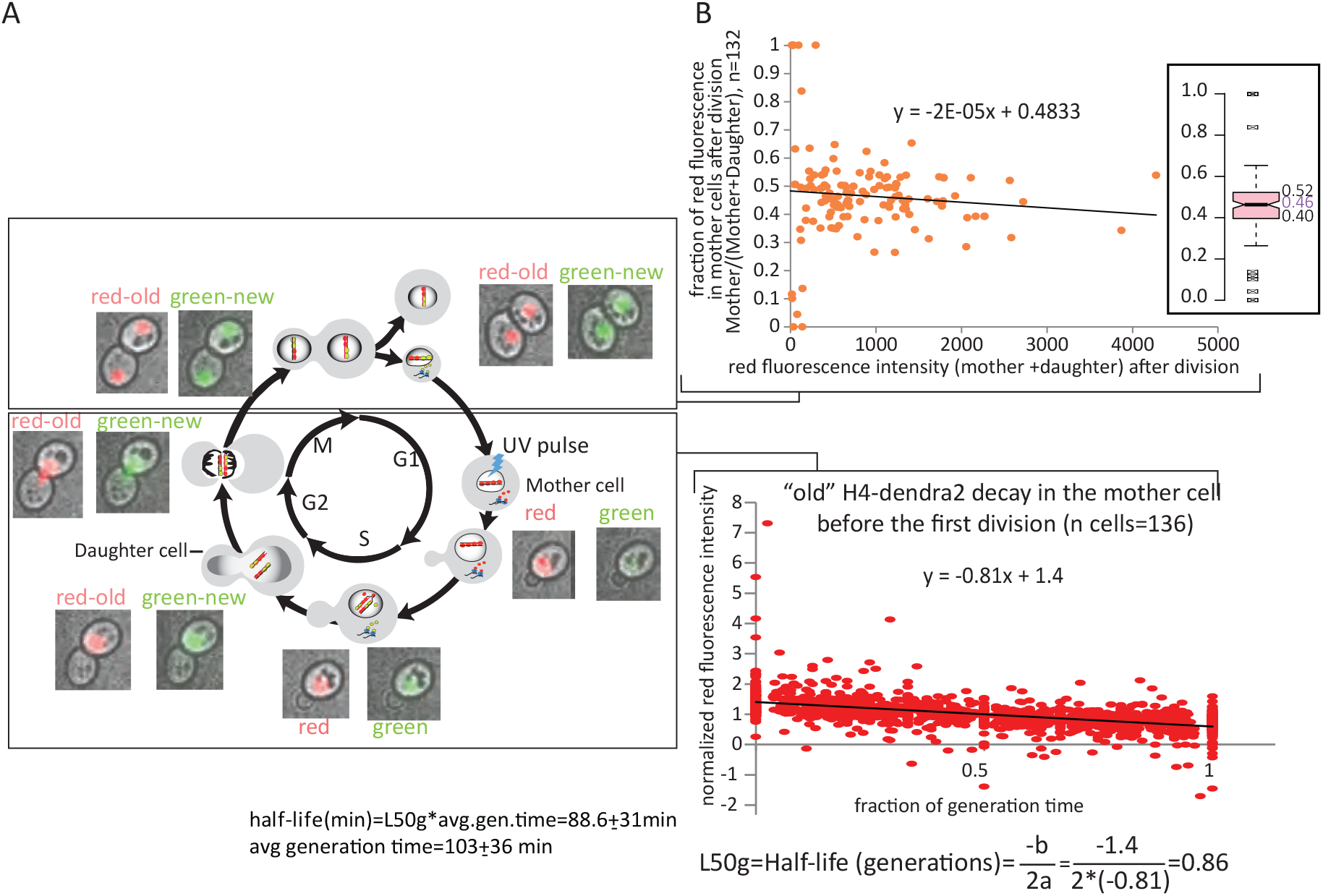
Determination of the half-life and maternal protein segregation pattern for H4-Dendra2. A. Old-red and new-green histone H4 distribution in the mother and daughter cells during the yeast cell cycle followed by HiLo live fluorescence microscopy. Cells were illuminated with UV light in G1 and images were taken every 6.5 min for six hours. Only the first cycle after photo conversion is shown. B. Half-life (bottom) and old/maternal H4-dendra2 partition (top) between mother and daughter cells. The x-axes represent all the time points taken before the first cell division after photo conversion from 136 mother cells calculated as fractions of generation time for each mother cell (bottom) and the sum of average red fluorescence intensities in the mother and her first daughter produced after photo conversion over the duration of the mother’s subsequent cell cycle (i.e. before the production of the second daughter after photo conversion) (top). The average generation time is an average of up to 3 cell cycle lengths of 136 mother cells. The red fluorescence intensities in both graphs have been subtracted from background fluorescence. The signal in the Y-axis of the bottom graph has also been normalized to the average fluorescence intensity from the time of photo conversion to the time of the first cell division for each mother cell. The average fraction of maternal H4 retained in the mother cell after cell division is estimated from the Y-axis cutoff in the top graph (0.48) and the median value in the box plot (top inset): 0.46.

The decrease in the amount of maternal proteins in the mother cell over time is determined from the decay rate of red fluorescence in the mother cell after photo conversion and before the first cell division. The measured rate of decay is used to calculate the half-life of the dendra2 fusion protein. Photo-bleaching did not affect the decay rate (**Supplementary Figure S1)**. The half-life of red fluorescence was calculated as described in **Figure 2B** and in **Materials and Methods**. The currently used methods for measuring protein half-lives are based on pulse-chase time-courses with labeled amino-acids in cell populations and require either careful calibration of total protein content between different time points or theoretical approximations of protein synthesis rates as in the classic [^35^S]-methionine and [^35^S]-cysteine pulse chase experiments^12^ or SiLAC MS experiments^13,14^, respectively. These experimental drawbacks are circumvented with our method because it is based on observations of single cells in vivo and it allows for direct and independent measurements of protein decay and net synthesis rates in each cell.

### Low abundance proteins and metabolic gene regulators Rpd3L, Fpr4 and Tup1 are retained in the mother

Half-lives and segregation patterns of maternal proteins for all 18 dendra2 fusions are summarized in **Figure 3**. Our analysis reveals an inverse correlation between protein retention in the mother cell and protein abundance. If protein partitioning between the mother and the daughter were purely stochastic, we expect a normal distribution of protein fractions retained in the mother with a mean ~0.5 and a variance that is inversely proportional to protein abundance. Indeed we observe that highly abundant proteins (estimated at >25000/cell, see Materials and Methods) are divided between the mother and the daughter into two equal sets in most cells as shown for histone proteins and Kap123 (nuclear transporter of histone proteins). Moderately abundant proteins (between 3000 and 6000 per cell) also segregate equally on average in the cell population but with bigger variability between cells, i.e. there is a higher fraction of mother or daughter cells in the population that have inherited disproportionally more maternal proteins. Moreover, if protein partitioning is random, the cells that inherit more proteins should be mothers or daughters in equal proportions, as we see for Rap1 (general transcription factor also involved in heterochromatin formation), Cbf1 (associates with centromeric DNA elements and kinetochore proteins) and Ioc3 (subunit of the nucleosome remodeler Isw1a). Finally, following the same reasoning, small quantities of proteins (less than 1300/cell) will segregate asymmetrically in an even larger fraction of cells, ending up in mothers and daughters in equal proportions, than moderately abundant proteins but the mean of the protein fraction retained in the mother should still be ~0.5. Surprisingly, this behavior is observed only for Sir2 (subunit of the heterochromatic Sir complex) and the linker histone H1. The other low abundance proteins Sin3, Vps75, Chd1, Asf1, Set2 and Hda2 are all preferentially retained in the mother, as are moderately abundant proteins: Rxt3, Fpr4 and Tup1 (**Figure 3A, C**).

**Figure 3:**
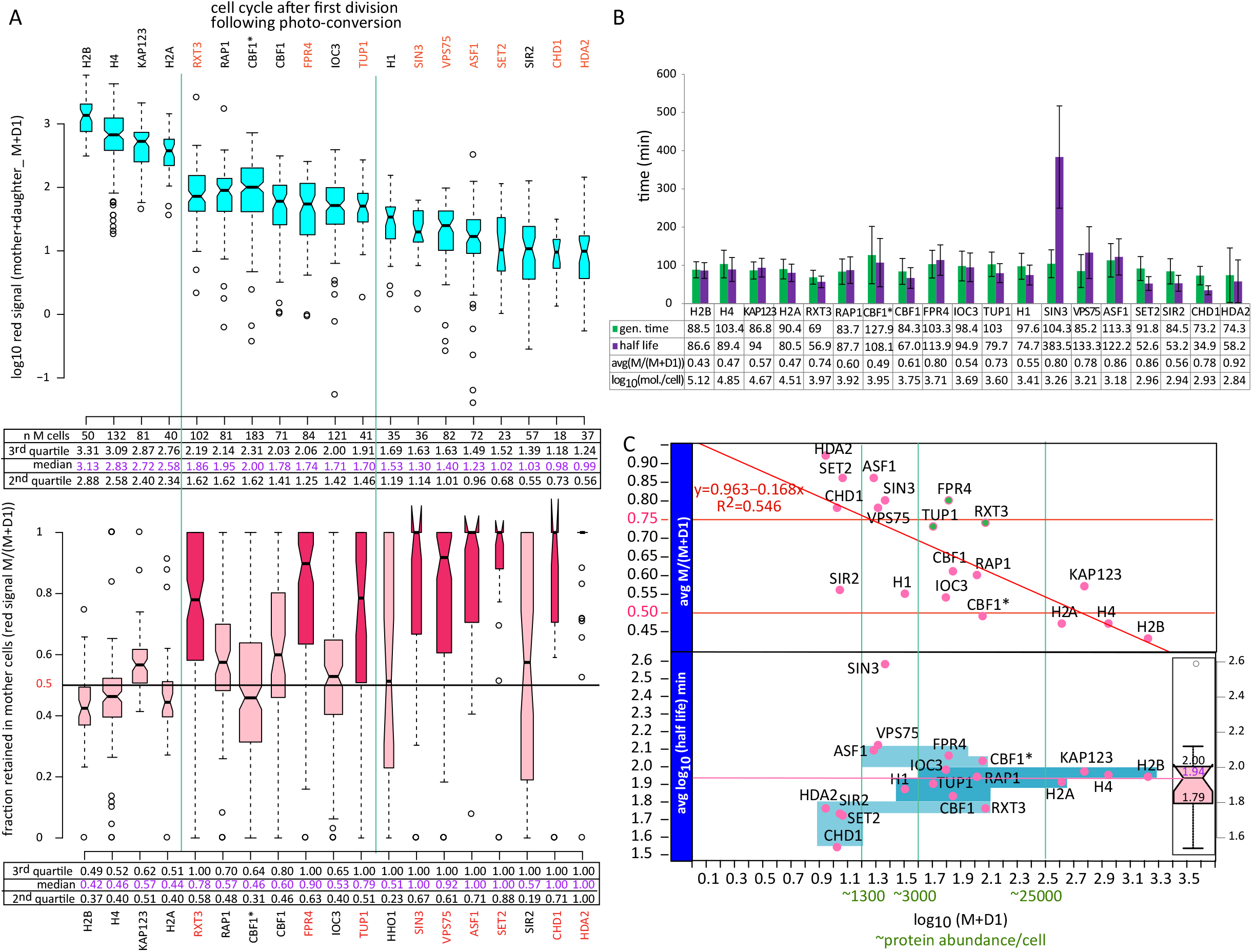
Segregation of 18 maternal chromatin proteins between mother and daughter cells after mitosis. A. Box plot distributions of average red fluorescence intensities (calculated as described in Figure 2B) from the mother M and her first daughter cell D1 after photo conversion (the number of analyzed mother/daughter pairs is indicated in the table below the top graph) (top) and the fraction of the total red fluorescence intensity shown on top that is retained in the mother M/(M+D1) (bottom). The tables below the graphs show the median values of the entire distribution (magenta) and the medians of the 2nd and 3rd quartile for each dendra2 protein fusion. The measured M and D1 values for IOC3 and SIR2 have been divided by 2 to correct for the double dendra2 tag of these two constructs. CBF1* is the strain with a lysD background that was used in the half-life measurements by mass spectrometry (Figure S2). B. Bar graph showing average cell generation times and half-lives of indicated dendra2 fusion proteins. Generation times were averaged over 2 to 3 generations in mother cells (number of cells indicated in A). Half-lives were determined from red fluorescence decay during the first cell cycle after photo conversion in mother cells and calculated as fractions of generation time as described in 2B and subsequently converted to minutes using the average generation time shown in the graph. The error bars represent the standard deviation from the average generation time. Maternal protein abundance (PA, sea-green x-axis in C) is proportional to the average total red fluorescence in the mother and daughter cells after mitosis (M+D1) for all measured mother-daughter pairs: PA=a*(M+D1). a=70000/((M+D1), H4)). 70000 is the estimated number of dendra2-histone fusion proteins bound to chromatin per cell: number of nucleosomes=70000=((genome size(12000000bp))/(nucleosome footprint +linker (160bp)))- (nucleosome free regions(5000)), only one histone gene of the two copies in the yeast genome has the dendra2 fusion). The efficiency of photo-conversion (ranging from 50 to 90%, data not shown) has not been taken into account in the calculation since conversion efficiencies varie from cell to cell and only an estimate of the order of magnitude of protein abundance is sufficient for the analysis. C. Correlation between protein abundance and half-life (bottom panel) and retention in the mother cell after mitosis (top panel). Bottom panel: The four blue rectangles (bottom to top) encompass proteins from the 1st to the 4th quartile from the box plot distribution of half-lives on the right, respectively. Low abundance proteins (<1300/cell) tend to be in the 1st quartile and high and moderately abundant proteins (>3000/cell) tend to be in the 3rd and 4th quartiles, with the exception of SIN3 (a low abundance protein with a long half-life). Top panel: Highly (>25000/cell) and moderately (between 3000 and 10000/cell) abundant maternal proteins are distributed stochastically between the mother and the daughter during mitosis, with a mean retention in the mother around 50% and variance inversely proportional to protein abundance (as shown in A). RXT3, FPR4 and TUP1 with a high retention bias in the mother are the exception. Low abundance proteins (between 600 and 3000/cell) have a clear retention bias for the mother cell with the exception of H1 and SIR2 which are distributed stochastically similar to highly abundant proteins.

### Half-life durations of abundant proteins are equal to cell cycle length

Our measurements of decay rates of red fluorescence also revealed that the half-lives of the examined proteins are comparable to cellular generation times except for low abundance proteins whose half-lives are ~50% of generation time (**Figure 3B-C**). This means that highly and moderately abundant proteins turn over completely in two generations. These proteins could therefore potentially transmit epigenetic information from one cell generation to the next if they stay associated with the relevant genomic loci after DNA replication and mitosis. The case for epigenetic inheritance of low abundance proteins is somewhat more difficult to make. Since these proteins turnover within one cell generation, epigenetic information could only be transmitted if newly synthesized proteins were exchanged with the old proteins directly on chromatin in order to preserve and transmit the information on their underlying genomic locations.

Curiously, Sin3 (component of Rpd3S/L histone deacetylase complexes) appears to have a half-life that is almost four times longer than the cell generation time. While the significance of this result is not clear it is also the only half-life value that matches half-life estimates measured by pulse^15^. The other half lives are on average 6 times shorter than the MS measurements (**Figure S2**). One possible explanation for this discrepancy is that the dendra2 tag destabilizes the protein. The results of a mass spectrometry SiLAC assay of cell extracts from two dendra2 strains (CBF1dendra2 and H1dendra2) from a time course after switch from Heavy Lysine to Light Lysine (**Figure S2B**) show that the tagged and untagged proteins have similar half-lives suggesting that dendra2 has no effect on protein stability. Moreover, our half-life estimates from the SiLAC experiment are closer to our halflife measurements from dendra2 fluorescence decay (**Figure S2C**) than to the published values mentioned above^15^. The discrepancy is therefore not due to the technical differences or experimental conditions in either experimental method used to measure half-lives.

The difference stems instead from the calculations used to derive half-lives in^15^ and our own calculations. Unlike in our assay where the protein decay rate is measured directly from the decay rate of red fluorescence in each cell, in SILAC-LC-MS/MS the protein decay rate is indirectly estimated from the decrease in the H/L ratios (**Figure S4 and Materials and Methods**) in a cell population over time after the switch to “light” medium. It is consequently necessary to make several assumptions in order to estimate protein decay rates and half-lives. First, it is generally assumed that protein decay is an exponential process^13,15^ although that seems not to be the case for some proteins^16^. Second, one needs to make a guess on protein synthesis rates, which are not measured in the MS assay. The calculations in^15^ were based on half-life measurements in mammalian cells from^13^ where it was assumed that total protein amounts double every cell generation. This assumption has been confirmed for 40 proteins in mouse ES cells^17^ but does not seem to hold for yeast. According to our direct measurements of net protein accumulation rates described below, the cellular content for 13 out of the 18 proteins we measured does not double within one cell cycle (**Figure S3 C**). As a consequence of that assumption, half-lives of proteins with actual synthesis rates higher than the two fold increase in one cell generation will be underestimated, while the half-lives of proteins with low synthesis rates will be overestimated as shown in Figure S2C. In order to be able to compare our half-life values with the published dataset, we nevertheless used the same approximations in our calculations. Surprisingly, we obtain values that are at odds with the ones in^15^. The major difference in our calculations and the ones in^15^ is in the use of a correction for protein dilution due to cell division applied in the latter. Maternal protein amounts are reduced two fold every cell generation because ~50% are passed on to the daughter in most cases (Figure 3), but since we use total cell extracts from a population of mothers and daughters in the SILAC experiment the only process that causes the decrease of “old/heavy” proteins in the cell population is protein degradation. The correction used in Christiano et al (2014) is therefore not necessary and is actually the cause for the overestimation of protein half-lives in that study.

### Coordinated protein synthesis and degradation

Since the intensity of green fluorescence after photo conversion is proportional to the amount of newly synthesized protein, we were able to derive protein synthesis rates from the rates of increase in green fluorescence during the first cycle after photo conversion (**Figure S3 and Materials and Methods**). We also determined the distribution of green fluorescence in the mother and the daughter after cell division as we have done for red fluorescence. The fraction of green fluorescence in the mother cell is however less informative about protein inheritance patterns since the amount of green proteins in the daughter is the sum of proteins inherited from the mother and new proteins that are synthesized in the daughter. Interestingly, even with this limitation, we can still detect the preferential retention of Rxt3 and low abundance proteins Set2, HDA2, Asf1 and Vps75 in the mother cell (**Figure 4A**). Thus, the tendency of low abundant proteins to stay in the mother cell is also evident in the “green” protein population (**Figure 4B**) that has not undergone photo-conversion, suggesting that the segregation pattern observed for “red” proteins is not due to damage potentially caused by the UV pulse.

**Figure 4:**
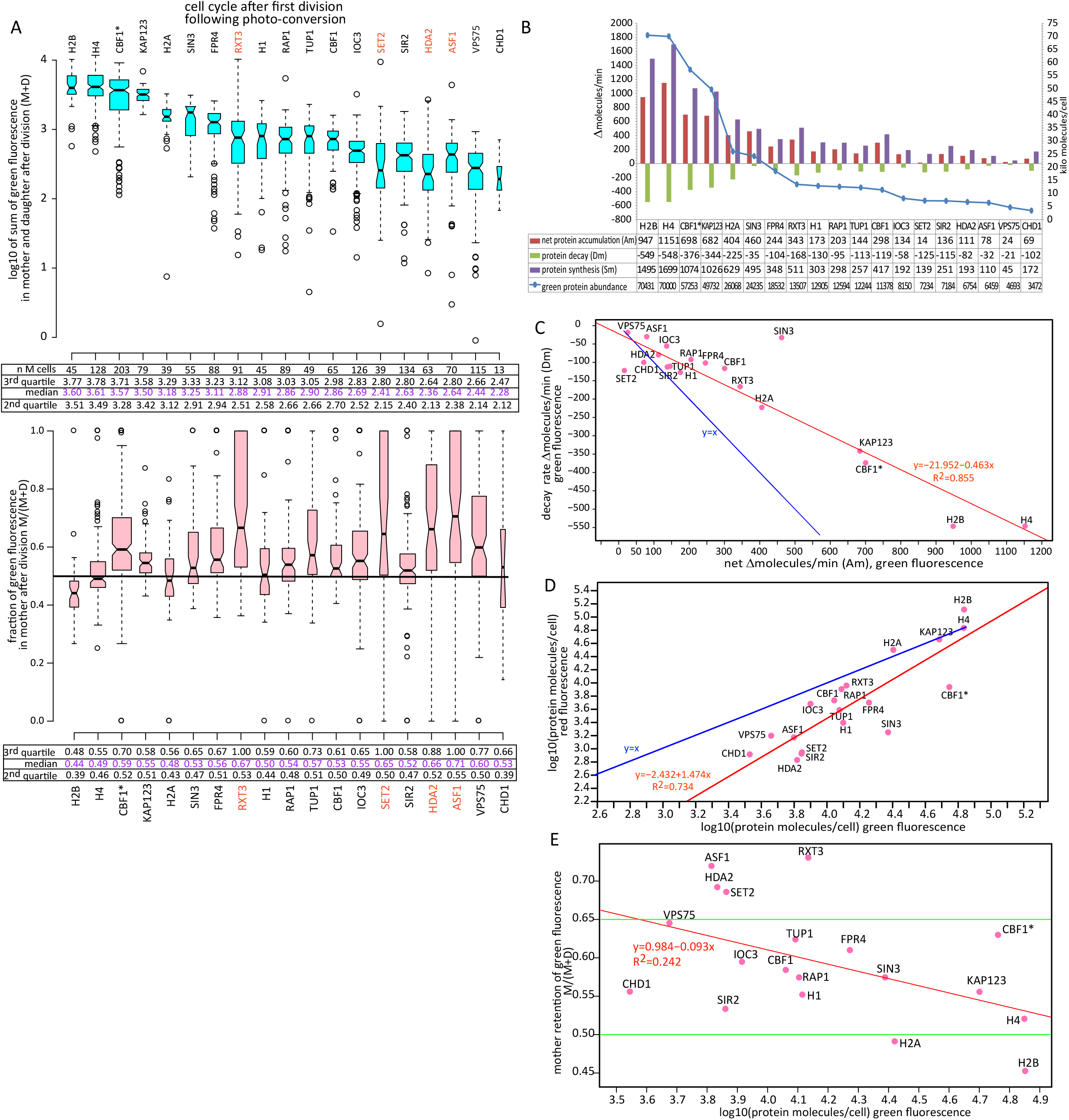
Segregation patterns between mother and daughter cells after mitosis and synthesis rates of 18 chromatin proteins. A. Box plot distributions of average green fluorescence intensities (calculated as described in Figure S2B) from the mother M and her first daughter cell D1 after photo conversion (the number of analyzed mother/daughter pairs is indicated in the table below the top graph) (top) and the fraction of the total green fluorescence intensity shown on top that is retained in the mother M/(M+D1) (bottom). The tables below the graphs show the median values of the entire distribution (magenta) and the medians of the 2nd and 3rd quartile for each dendra2 protein fusion. The measured (M+D) values for IOC3 and SIR2 have been divided by 2 in the top graph to correct for the double dendra2 tag of these two constructs. CBF1* is the strain with a lysD background that was used in the half-life measurements by mass spectrometry (Figure 5). B. Correlation between new (green) protein abundance per cell and green protein retention in the mother cell. The number of molecules per cell was estimated from the total green fluorescence in the mother and daughter cells after mitosis (M+D1): PA=a*(M+D1)/2. a=140000/((M+D1), H4). 140000 is the estimated number of dendra2-histone fusion proteins bound to chromatin in the mother and the daughter cells: number of nucleosomes per cell=70000=((genome size(12000000bp))/(nucleosome footprint +linker (160bp)))- (nucleosome free regions(5000)), only one histone gene of the two present in the yeast genome has the dendra2 fusion). Since most of these proteins are continuously synthesized we cannot distinguish between green proteins that were inherited from the mother and the ones that were newly synthesized in the daughter cell. Consequently most green proteins are equally distributed between the mother and the daughter except for ASF1, HDA2, SET2 and RXT3 which are retained in the mother as observed for maternal (old) proteins (Figure 3). The low abundance proteins have a tendency to stay in the mother cell as observed for “old” maternal proteins. C. Bar graph showing average net protein accumulation rates (Am), protein decays rates (Dm) and protein synthesis rates (Sm) in molecules/min calculated as described in Fig. S3B. Am, Dm, and Sm rates were converted from fold change/min (see Fig. S3B) to molecules/min by multiplying each rate with the corresponding PA (protein abundance per cell calculated as in B). D. Correlation between protein accumulation rates (Am) and protein decay rates in molecules/min. Values are based on green fluorescence intensity as in B-C. Synthesis and decay rates appear to be co-regulated with decay rates ~2 fold slower than protein accumulation rates (net protein production) (1/0.498=2.01), with the exception of SIN3 (net synthesis 13 fold faster than decay) and SET2 (decay is 8 fold faster than net synthesis).E. Correlation between old (red) and new (green) protein abundance (PA) per cell (log 10(protein molecules/cell)) calculated as in B. Due to faster synthesis rates compared to decay rates there is consistently more new “green” protein than old “red” protein in the cell. Note that the equal numbers of old and new histone is a consequence of normalization of all fluorescence intensities to the fluorescence intensity of H4 and the assumption that nucleosomes contain equal amounts of old and new histones as explained in B.

Our independent measurements of protein synthesis (using the population of “green” proteins) and decay (using the population of “red” proteins) have revealed an unexpected coupling of the two processes (**Figure 4D**). Surprisingly, protein degradation is correlated with protein synthesis with a net protein accumulation rate at ~2 fold the decay rate for most proteins we tested.

### Discussion

We have developed a live cell imaging method for quantitative measurements of protein behavior in single cells that allowed us to directly measure degradation and net synthesis rates as well as inheritance patterns of 18 chromatin proteins. The dendra2 tag and the UV pulse for photoconversion do not significantly affect protein stability as shown by similar protein half-lives measured by microscopy and SILAC-LC-MS/MS.

Our results show that “old” low abundance proteins have a tendency to remain in the mother cell after mitosis. The biological significance of this observation still remains to be explored but we speculate that the retention in the mother cell may be related to the potentially higher levels of oxidative damage in these proteins, which may also be the reason for their shorter half-lives. These proteins may have reached the end of their useful lives and are not being transferred to the daughter cell because their function has completely deteriorated by the time of cell division.

The retention of maternal Fpr4, Rxt3 and Tup1 in the mother cell is especially interesting, since all three are moderately abundant proteins that have been implicated in the regulation of transcription of inducible genes from metabolic pathways^18–20^. Fpr4 is a proline isomerase specific for P38 on histone H3. The activity of Fpr4 inhibits methylation of H3K36 at promoters, which stimulates the induction of MET16 and HIS4^18^. Tup1 and Rxt3 have both been implicated in the transcriptional memory of genes that are induced or repressed during carbon source shifts, respectively. Rxt3 is a subunit of the histone deacetylase Rpd3L, which is recruited to promoters of genes that need to be repressed when galactose is the carbon source. It has been found recently that Rpd3L is necessary for transcriptional memory of repression i.e. for faster repression rates upon repeated exposure to galactose during carbon source shifts^19^. Tup1 on the other hand appears to be necessary for long term transcriptional memory and faster reactivation of galactose inducible GAL genes^20^. Although transcriptional memory seems to be preserved for several cell generations, the above studies have not addressed how these factors partition between the mother and the daughter and consequently whether mothers transmit the information to their daughters. Since our assay was done in constant carbon source conditions (cells grew in glucose before and during imaging) the preferential retention of these factors in the mother cell may not be connected to transcriptional memory of genes involved in carbon source metabolism unless mother retention is the default state in a constant carbon source environment. An intriguing possibility is that these factors may be involved in the transcriptional memory of stress response genes that were induced in the mother cell during photo-conversion and that the memory of stress response activation stays in the mother. Future studies should elucidate whether these proteins remain associated with relevant genomic loci throughout cell division, which is a prerequisite for epigenetic inheritance.

Our analysis of protein decay and synthesis rates has also revealed some unexpected features of budding yeast protein biology. It is remarkable that even though protein synthesis and degradation are distinct and seemingly independent processes, they are precisely coordinated. Protein homeostasis with equal synthesis and decay rates is not achieved as might be expected. The higher synthesis rate results instead in a steady supply of new proteins. In other words, the level of chromatin proteins steadily increases throughout the cell cycle and never reaches a steady-state. Consequently, at any given time during the cell cycle newly synthesized “younger” proteins will represent the majority of the total protein population as shown in **Figure 4E**. Since old proteins are more likely to be damaged, the accumulation of new proteins may be an adaptation to ensure that the cell has optimal amounts of functional proteins at its disposal. Since only about a third of synthesized proteins whose half-life lasts one cell cycle are degraded in one cell generation, the cell has to rely on two processes to maintain optimal protein levels: protein degradation and dilution through cell division. The accumulation of new proteins may also be a hallmark of asymmetrically dividing cells in which the mother cell has to supply all the proteins necessary for her daughter’s initial growth until the daughter cell has produced enough proteins on her own.

Measurements of a larger protein set should confirm whether the processes we uncovered such as the coupling of protein synthesis and degradation that favor the accumulation of new proteins in the cell, half-life duration of one cell cycle and the stochastic and symmetric partitioning of proteins between the mother and its daughter, represent general trends in protein biology of asymmetrically dividing cells. Likewise, further studies of the mechanisms responsible for the asymmetric repartition of Rxt3, Tup1 and Fpr4 proteins should help us better understand the potential role of these proteins in epigenetic inheritance of transcriptional memory.

## Materials and Methods

### Yeast Strains and Dendra2 Plasmid Construction

The plasmid pDendra2NatMX was constructed by ligation of the NheI-HpaI Dendra2 fragment from pDendra2-C (Clontech) and the PvuII linearized pAG25 vector (Addgene). The Dendra2 restriction fragment was blunt-ended using End-it Repair (Epicenter) before ligation. The correct orientation of Dendra2 relative to the NatMx marker was verified by sequencing. The pDendra22xNatMX plasmid with two Dendra2 tags in tandem repeat was constructed as above with serial cutting and pasting of Dendra2 into the pAG25 vector using NdeI and MfeI restriction enzymes for the first insertion and NheI and AflII for the second. Appropriate restriction sites were introduced into the Dendra2 insert during PCR. (primer pair for first insertion: 5’CATATGGGTGCTGCTAGCGGTGCTCTTAAGAACACCCCGGGAATTAACCT 3’CAATTGGTCATAGCTGTTTCCTCTTATCTAGATCCGGTGGATC Primer pair for second insertion: 5’GCTAGCAACACCCCGGGAATTAACCT 5’CTTAAGGGAGCAGGTGCTGGTGCTGGTGCTGGAGCA TCTAGATCCGGTGGATCCCG).

Dendra2 was then fused to the C-terminus of chromatin proteins of interest by homologous recombination into the JOY1 parent strain (MATa ura3D leu2D his3D met15D bar1D::HIS5, BY4741 background) and integration was verified by PCR (resulting genotype: MATa ura3D leu2D his3D met15D bar1D::HIS5 geneX-Dendra2: NatMX). Integration and verification PCR primers are listed in Table S1.

For the Dendra2 strains for mass spectrometry measurements, Dendra2 was fused to HHO1 or CBF1 in the lys2 deletion mutant from the barcoded YSC1053_KO deletion library (GE Healthcare/dharmacon) (MATa ura3D leu2D his3D1 met15D lys2D::KanR).

Since N and C terminal tagging inactivates Rap1, dendra2 was inserted within the N-terminal domain of Rap1 in place of the GFP tag used in^21^. GFP in the pAH52 plasmid obtained from A. Taddei has been replaced by dendra2 using the SLiCE method^22^ and the Pst1 linearized plasmid with Dendra2-Rap1 has then been recombined with the genomic Rap1 gene.

### Yeast cell culture for live cell imaging

Cells were grown at 30°C in 3ml of SCD (Synthetic Complete Dextrose) medium to OD ~ 0.5, and were then concentrated by centrifugation. 3μl of the cell pellet was injected under the 0.8% agarose/SCD layer that had been poured into each well of an 8-well glass bottom microscopy plate (BioValley), in order to screen eight different strains in each time course.

### Yeast cell culture and protein cell extract preparation for mass spectrometry- SILAC- LC-MS/MS

Cells were grown in SCD-Lys medium supplemented with heavy Lysine (L-Lysine-2HCL (13C6 99%, 15N2 99%) (Cambridge Isotope Laboratories, Inc.) (20mg/L) at 30°C for 48hrs (the OD was kept at 0.5). The culture was switched to light lysine and aliquots were taken at 30, 60, 90, 120, 150 and 180min. Total cell protein extracts were purified as follows. 8ml of 100% TCA was added to 40ml cell culture aliquots, and cells were pelleted after 10min incubation on ice and pellets were washed and re-supended in 300μl cold 10% TCA. Cells in the TCA suspension were mechanically spheroplasted by bead beating with 0.5mm Zirconium Sillicate beads in a bullet blender (Next Advance) for 4 times x 3 min (intensity 8) at 4°C. The spheroplast pellets were then re-suspended in 2xSDS-PAGE loading buffer (4% SDS, 100mM Tris pH=6.8, 20% glycerol, 200mM DTT) and TCA was neutralized with 1M Tris-HCl pH 8.7. Cell debris was pelleted (5min, 17000g) after a 5 min incubation at 95°C, and the supernatant with the protein cell extract was used for mass spectrometry.

### SILAC- LC-MS/MS analysis

Proteins from cell extracts were separated by SDS-PAGE. Proteins from 2 different gel fractions containing the dendra2 tagged and untagged versions of Cbf1 and H1 proteins were subjected to LysC (ThermoScientific) digestion. Obtained peptides were analysed online using Qexactive HF mass spectrometer (ThermoScientific) coupled with an Ultimate 3000-RSLC (ThermoScientific) fitted with a stainless steel emitter (ThermoScientific). Desalting and preconcentration of samples were done on-line on a Pepmap^®^ precolumn (0.3 mm x 10 mm, ThermoScientific). A gradient consisting of 0-40% B in 120 mn, 90% B during 5 min (A = 0.1% formic acid, 2% acetonitrile in water; B = 0.1 % formic acid in acetonitrile) at 300 nl/min was used to elute peptides from the capillary (0.075 mm x 150 mm) reverse-phase column (Pepmap^®^, ThermoScientific). Spectra were recorded using Xcalibur software (v 4, ThermoScientific) and acquired with the instrument operating in the information dependant acquisition mode throughout the HPLC gradient. A cycle of one full-scan mass spectrum (375–1,500 m/z) at a resolution of 60,000 (at 200 m/z), followed by 12 data-dependent MS/MS spectra (at a resolution of 30,000, isolation window 1.2 m/z) was repeated continuously throughout the nanoLC separation. Analysis was performed using Maxquant software^23^. All MS/MS spectra were searched using Andromeda^24^ against a decoy database consisting in a combination of yeast entries from Reference proteome database (release 2018_04, https://www.uniprot.org/) and 250 classical contaminants, containing forward and reverse entries. Default SILAC search parameters were used. Briefly, first search precursor mass tolerance was set to 20 ppm, and main search (after recalibration) to 6 ppm. A maximum of 2 mis-cleavages was allowed. Search was performed allowing variable modifications: Oxidation (Met), Acetylation (N-term) and with one fixed modification: Carbamidomethyl (Cys). FDR was set to 0.01 for peptide and proteins, and minimal peptide length to 7. Re-quantify option was used to perform ratio calculation.

#### Half-life calculations from SILAC- LC-MS/MS

Normalized ratios r=H/L are obtained by dividing the average H/L ratio of each protein at every time point with the average H/L ratio for that protein at t=30min, which was considered as time 0.The t=0min time point was discarded because the linear fit to the equation (4) was better without this point. Heavy protein *H* decay over time *t* is assumed to be exponential: (1) *H*(*t*) = *P*_o_*e^−kdt^*; *P_0_* is the total protein at time 0 and is assumed to be equal to the normalized fraction of heavy protein *H_0_* at time 0: *P_0_=H_0_=1*; kd is the decay rate. Total protein *T* content is assumed to double within one cell generation: (2) 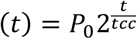; t_cc_ is the cell cycle length and was estimated from OD measurements over the time course. The change in light protein *L* over time is described with the function: (3) *L(t)=T(t)-H(t)*. The change in r over time can be plotted as ln (1+1/r) versus t according to the equation (4) below derived from the above functions (1), (2) and (3):

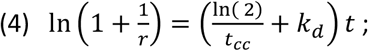

The decay rate is calculated from the slope of the linear fit to the scatter plot above: 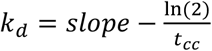. Finally protein half-life t_1/2_ is calculated from the decay rate: 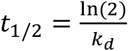

### Microscopy

We used a wide field inverted microscope for epifluorescence and TIRF acquisition (Nikon) under the HiLo setting, with a 60X water objective with a water dispenser, and a EMCCD Evolve 512 Photometrics camera (512*512, 16μm pixel size). The time courses on growing cells were performed at 30°C. Photo-conversion was done with a Lumencor LED lamp and the DAPI filter. Green and Red fluorescence was detected with 488nm (100mW at 360 μW) and 561nm (100mW at 300μW) lasers for excitation and GFP and TAMRA filters for emission, respectively. The Neutral Density ND4 or ND8 filters were used during photo-conversion with the 390nm LED lamp at 60% capacity for 1min. Pictures in bright field, and with the red and green lasers (Exposure time= 300ms, EM Gain= 1000 and the Hilo angle= 62°) were taken every 6.5min for 6.5hrs or as indicated. We used the maximum Intensity from 5 z-stacks with a 1μm gap for the analysis of fluorescent images.

### Image and Data analysis

ImageJ (Wayne Rasband, NIH, USA) and its plugin BudJ (Martí Aldea, Institut de Biologia Molecular de Barcelona) were used for image analysis. BudJ tracks marked cells throughout the time course and records their fluorescence intensities in each time frame The data output from BudJ was further processed using custom Perl and R scripts (available upon request).

#### Half-Life calculation

The background signal for each mother cell defined as the average fluorescence intensity after the inflection point of the time course (i.e. after the red signal has stopped decreasing and has flat lined, the time course typically spans three to four cell divisions), has been subtracted from the fluorescence intensity at each time point. All intensities from time points in one generation were then normalized to the average signal for that generation. Time points were converted from minutes to fractions of generation time (i.e. the time it took for the mother cell to produce its first daughter after photo conversion) for each mother cell in order to eliminate the variability in generation times between different mother cells. Normalized first generation intensities of all mother cells were then grouped in a common scatter plot and the half-life of the red fluorescence signal was calculated from the slope of the linear fit (Figure 2B bottom):

Half-life in generations (L50g):

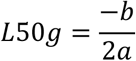

Where *b* is the y axis cut-off and *a* is the slope. The half-life in min is calculated from L50min=L50g*(average generation time). The average generation time is the average doubling time in minutes for all recorded cell divisions of all the observed mother cells.

#### Calculation of the maternal and new protein fractions retained in the mother cell

The fraction of proteins in the mother cell after cell division is calculated from Mf/(Mf+D1f), where Mf is the average fluorescence intensity in the mother cell M from the time when nuclei are fully partitioned between the mother (M) and its first daughter after photo conversion (D1) up to the appearance of the bud for the second daughter (D2). D1f is the average fluorescence intensity in the first daughter D1 during the same time interval used to calculate Mf.

#### Protein synthesis rate and protein abundance calculation

The background signal for each mother cell defined as green fluorescence intensity immediately after photo-conversion has been subtracted from the fluorescence intensity at each subsequent time point. Green fluorescence intensities at each time point in the first generation for each mother cell were normalized to the average fluorescence intensity from the time of photo conversion to the time of the first cell division as in half-life calculations for red fluorescence intensities. Time points were converted from minutes to fractions of generation time (i.e. the time it took for the mother cell to produce its first daughter after photo conversion) for each mother, as above for half-life calculations. Normalized first generation intensities of “new” green fluorescent fusion proteins in all mother cells were then grouped in a common scatter plot and the net rate of protein accumulation in fold increase per cell generation (Ag) was calculated from the slope of the linear fit (Figure S2B bottom). The protein synthesis rate Sg in fold increase per generation is then obtained from Sg=Ag-Dg, where Dg is the rate of protein decay in fold decrease per generation calculated as described above (Figure 2B bottom). Sg, Ag and Sg were converted from fold change/generation to fold change/min (Sm, Am and Dm respectively) by dividing each with the average generation time in min (Figure S2B).

The number of molecules of protein x per cell (PA_x_, protein x abundance) was estimated from total green (Figure 4) or red (Figure 3) fluorescence in the mother and daughter cells after mitosis (M+D1)_x_ (averaged over all measured mother daughter pairs) normalized to the estimated number of histone H4-dendra2 per cell as follows:

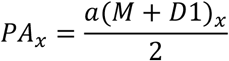

with

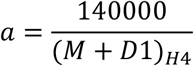

140000 is the estimated number of dendra2-histone fusion proteins bound to chromatin in the mother and the daughter cell: number of nucleosomes in one cell is 70000 =((genome size(12000000bp))/(nucleosome footprint +linker (160bp)))- (nucleosome free regions(5000)), since only one histone gene of the two copies in the yeast genome has the dendra2 fusion. (M+D1)_H4_ is the average total fluorescence of histone H4-dendra2 fusions. Am, Dm, and Sm rates were converted from fold change/min to Δmolecules/min by multiplying each rate with the corresponding PA (protein abundance per cell).

---

**Table S1: Yeast strains Insertion and verification PCR primers**

## Acknowledgments

PV constructed strains and plasmids, optimized and performed the live cell imaging experiments, and analyzed images. ST constructed strains and prepared samples for mass spectrometry, performed live cells imaging experiments and analyzed images with assistance from AF. FRC and IS constructed strains. MRL conceived and designed the experiments, wrote the manuscript and developed the Perl/R scripts for analysis. KK and SU performed the MS pulse SILAC analysis. We thank Virginie Georget (MRI, Biocampus, Montpellier) for her invaluable help with the microscope set-up. This study has been supported by the ANR 2014 GenChroSeg grant (MRL).

---

**Figure S1:**
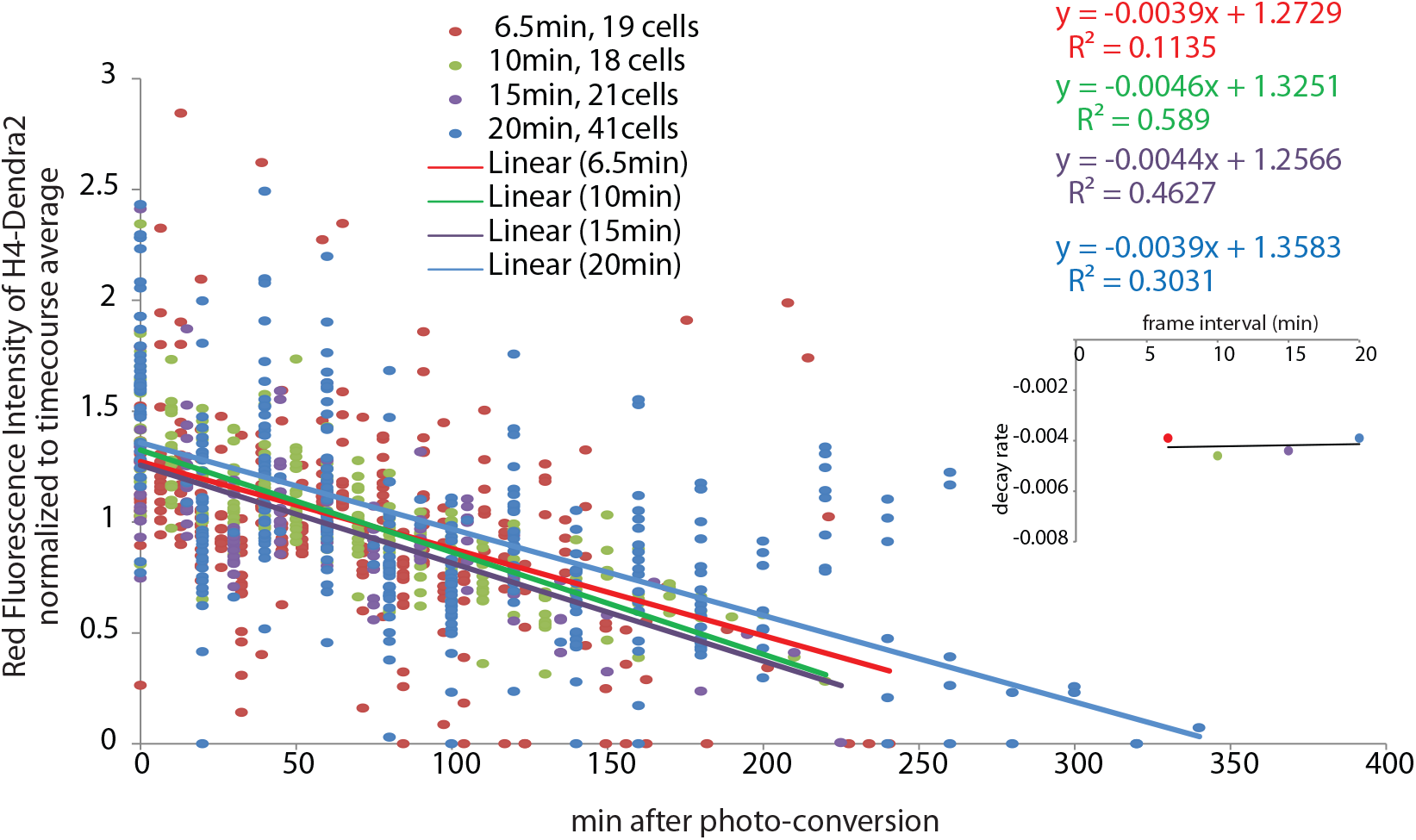
Test for photobleaching of red fluorescence. The intensity of red fluorescnce after photo conversion was recorded between the time of photo conversion and the first division of H4-Dendra2 mother cells. The red signal from the indicated number of mother cells was recorded at varying time intervals in four time courses, every: 6.5, 10, 15 or 20min. Since fluorescence decays at the same rate in all four time courses (as shown by the equations of the linear fit for each timecourse and the graph in the inset on the right; note the equations are of the same color as their corresponding line in the graph and figure legend), the observed decrease in red fluorescence is not due to photo-bleaching in our experimental conditions. Otherwise the decay rate would have been inversely proportional to the interval between frames (i.e more negative for the 6.5min interval than for the 20min interval) because cells would have been exposed to 15 laser pulses in 100min with the 6.5min interval and only 6 pulses in 100min with the 20min interval.

**Figure S2:**
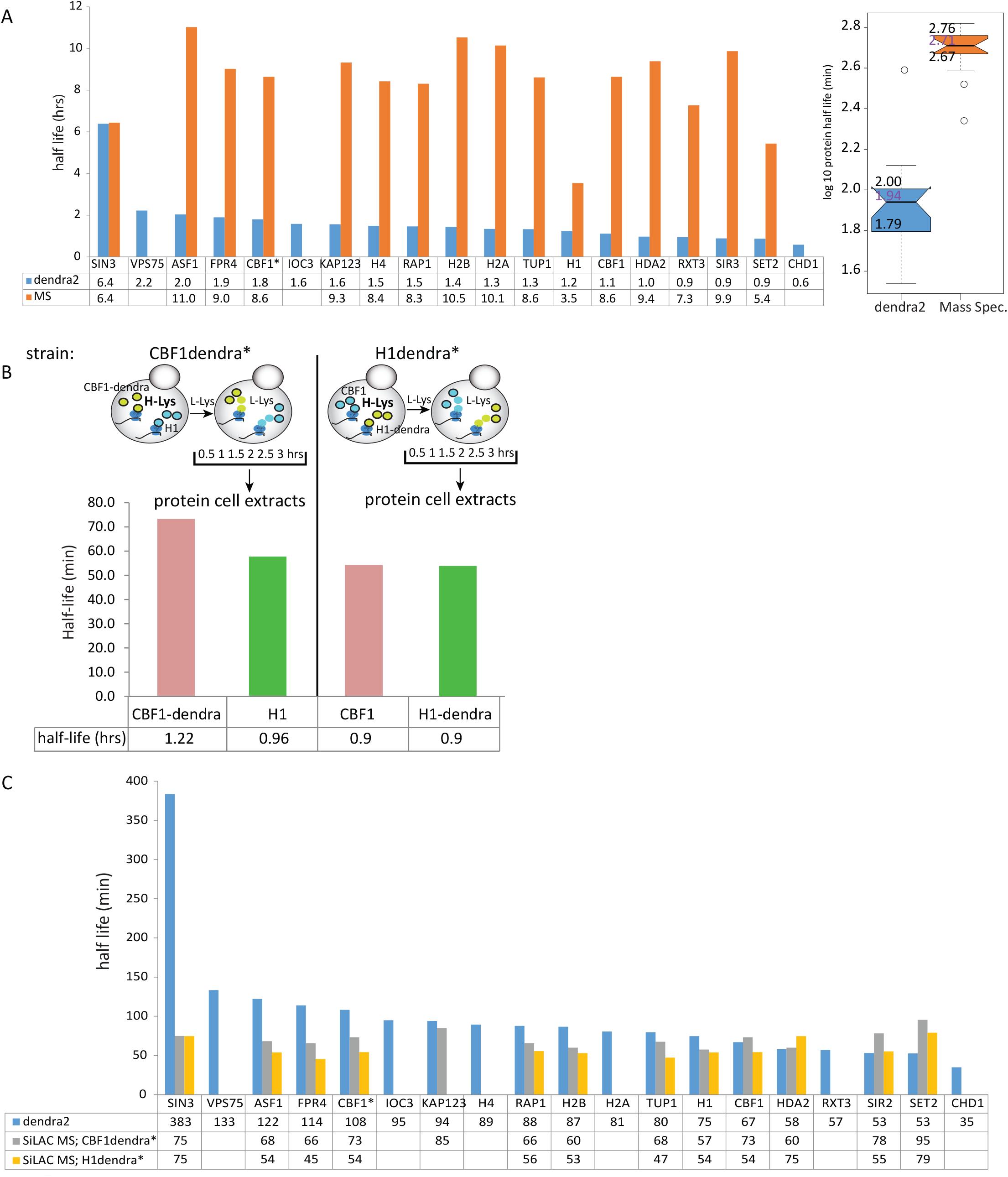
**A.** Comparison of half-lives estimates between dendra2 decay rates (Figure 3) and SiLAC-MS (Christiano et al., 2014). CBF1* is the lysΔ strain with CBF1dendra2 used in the SiLAC experiment in B. The box plot distributions of half lives in the bar plot (left) are shown on the right. **B.** SiLAC-MS in CBF1dendra2 and H1dendra2 strains. The two strains were grown in Heavy-lysine (H-Lys) and then switched to Light-lysine. Total cell extracts were collected after indicated times and the Heavy/Light peptide ratios for CBF1dendra2 and H1 (from the CBF1dendra* strain with lys2D), and CBF1 and H1dendra2 (from the H1dendra2*strain with lys2D) were calculated from Mass Spectrometry profiles. The half lives were calculated using the H/L ratios as described in Christiano et al. (2014) but without the correction for protein dilution by cell division. C. Comparison of half-lives between dendra2 decay from A and the SiLAC MS experiment from B (Half-lives calculated as in B). The generation times for the MS SiLAC experiment that were used for half-life calculations (Figure S4) were: CBF1dendra*: 273min and H1dendra*:286min.

**Figure S3:**
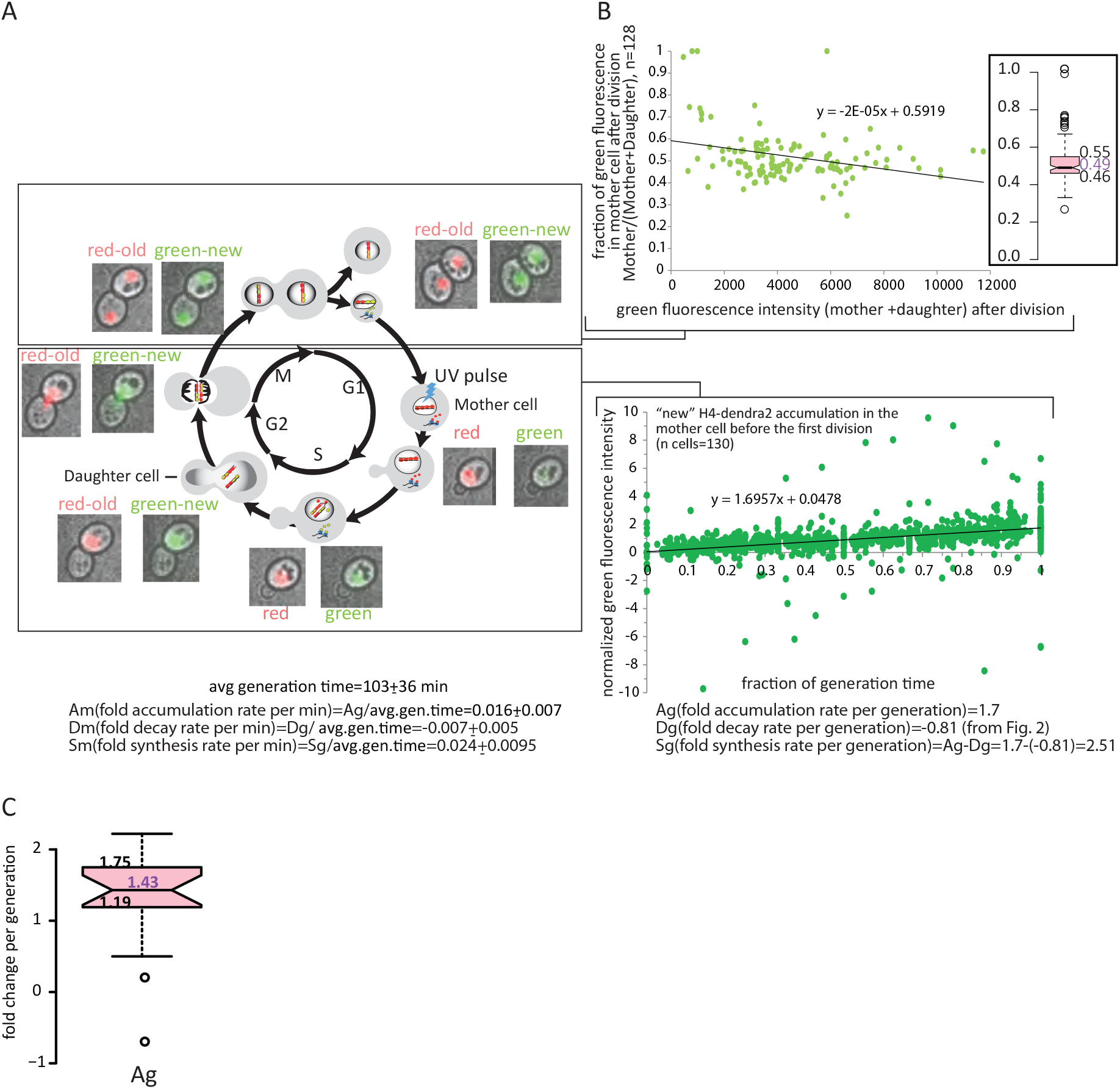
Determination of the new protein accumulation rate and segregation pattern for H4-Dendra2. **A.** Old-red and new-green histone H4 distribution in the mother and daughter cells during the yeast cell cycle followed by HiLo live fluorescence microscopy. Cells were illuminated with UV light in G1 and images were taken every 6.5 min for six hours. Only the first cycle after photo conversion is shown. **B.** Protein accumulation rate (bottom) and new H4-dendra2 partition (top) between mother and daughter cells. The x-axes represent all the time points taken before the first cell division after photo conversion from 130 mother cells calculated as fractions of generation time for each mother cell (bottom) and the sum of average green fluorescence intensities in the mother and her first daughter produced after photo conversion over the duration of the mother’s subsequent cell cycle (i.e. before the production of the second daughter after photo conversion) (top). The average generation time is an average of up to 3 cell cycle lengths of 130 mother cells. The green fluorescence intensities in both graphs have been subtracted from background fluorescence (i.e. the green fluorescence intensity at time 0 after photo-conversion). The signal in the Y-axis of the bottom graph has also been normalized to the average fluorescence intensity from the time of photo conversion to the time of the first cell division for each mother cell. The average fraction of maternal H4 retained in the mother cell after cell division is estimated from the average of the Y-axis cutoff in the top graph (0.59) and the median value in the box plot (top inset): 0.49: 0.54 **C.** Box plot distribution of Ag values for all 18 proteins from Figures 3 and 4 calculated as in B.

**Figure S4:**
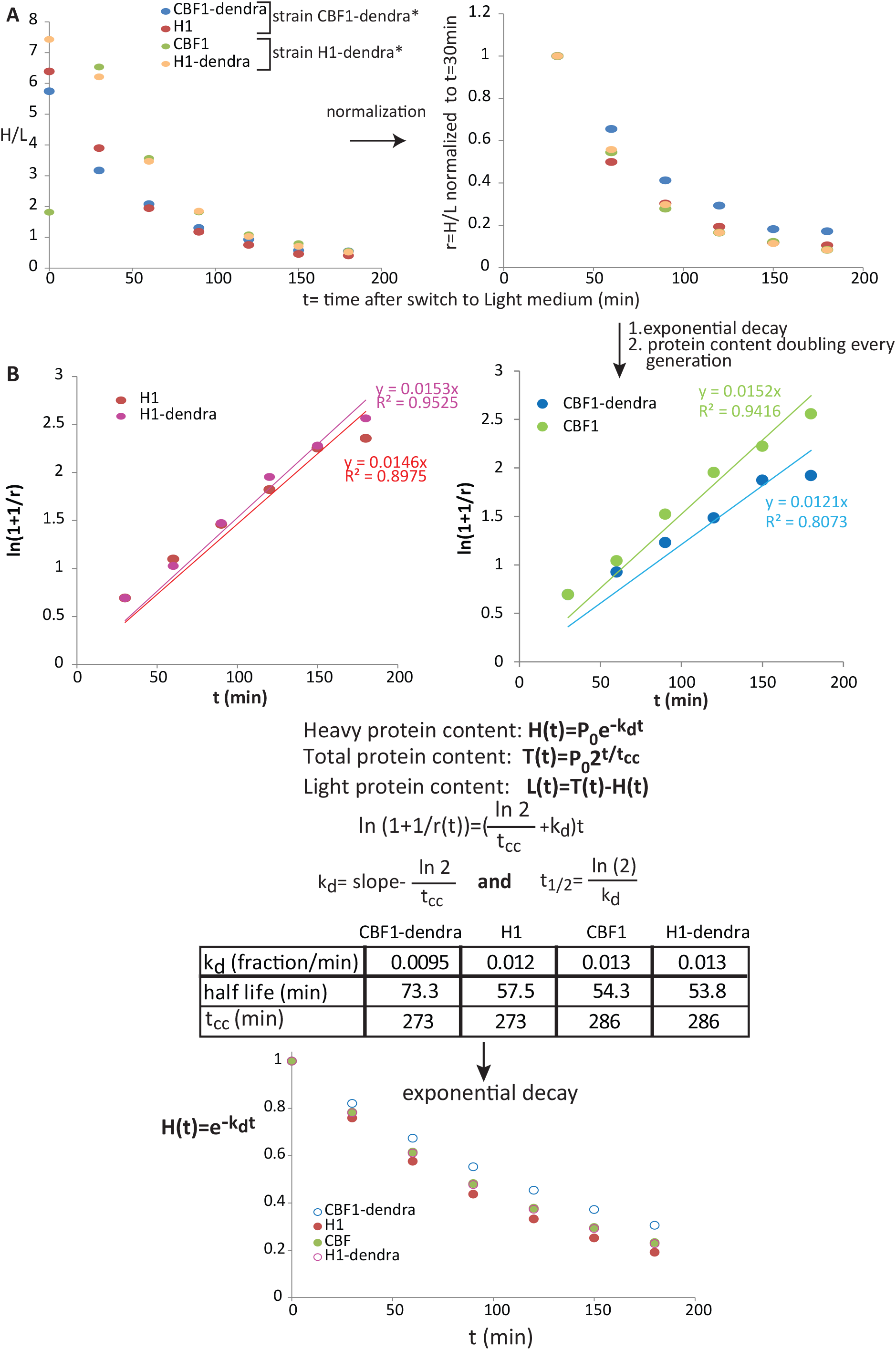
Half-life calculations from the SiLAC MS experiment in CBF1dendra*(lys2D) and H1dendra*(lys2D) **A.** Scatter plot of non-normalized H/L ratios (left) and r=H/L ratios normalized to t=30min (right), each H/L ratio is an average of 9 and 8 peptides for H1 and CBF1, respectively. t=0 is the time before switch to Light Lysine medium, and showed too much variability in H/L between peptides of the same protein and was not considered in the calculations. **B.** Half life calculations assuming exponential decay and a doubling of total protein every cell generation as in Schwanhausser et al. (2011). The linear equation for the plots in the top panel of ln(1+1/r) versus t is derived from the equations below the graph. P_0_-total protein, P_0_~=H_0_^=^100% at t=30min; t_cc_- generation time for the conditions used in the SiLAC time-course. The decay rate and half life were calculated using the linear fit curves shown in the top panel. The Y_intercept was set to 0 for the fit because y_intercept=0 in the derived equation. The bottom plot shows the exponential decay curve of heavy protein using the parameters calculated above.

